# The effect of anteroposterior perturbations on the control of the center of mass during treadmill walking

**DOI:** 10.1101/848853

**Authors:** Maud van den Bogaart, Sjoerd M. Bruijn, Jaap H. van Dieën, Pieter Meyns

## Abstract

Shifts of the center of pressure (CoP) through modulation of foot placement and ankle moments (CoP-mechanism) cause accelerations of the center of mass (CoM) that can be used to stabilize gait. An additional mechanism that can be used to stabilize gait, is the counter-rotation mechanism, i.e., changing the angular momentum of segments around the CoM to change the direction of the ground reaction force. The relative contribution of these mechanisms to the control of the CoM is unknown. Therefore, we aimed to determine the relative contribution of these mechanisms to control the CoM in the anteroposterior (AP) direction during a normal step and the first recovery step after perturbation in healthy adults. Nineteen healthy subjects walked on a split-belt treadmill and received unexpected belt acceleration perturbations of various magnitudes applied immediately after right heel-strike. Full-body kinematic and force plate data were obtained to calculate the contributions of the CoP-mechanism and the counter-rotation mechanism to control the CoM. We found that the CoP-mechanism contributed to corrections of the CoM acceleration after the AP perturbations, while the counter-rotation mechanism actually contributed to CoM acceleration in the direction of the perturbation, but only in the initial phases of the first step after the perturbation. The counter-rotation mechanism appeared to prevent interference with the gait pattern, rather than using it to control the CoM after the perturbation. Understanding the mechanisms used to stabilize gait may have implications for the design of therapeutic interventions that aim to decrease fall incidence.

**Summary statement:** Understanding the mechanisms used to stabilize gait during unperturbed and perturbed walking may have implications for the design of therapeutic interventions that aim to decrease fall incidence.

## Introduction

Stable gait, defined as gait that does not lead to falls (Bruijn et al., 2013), requires control of the position of the body center of mass (CoM) relative to the base of support (BoS, i.e. the area within an outline of all points on the body in contact with the support surface). In gait, the BoS is formed by those parts of the feet that are in contact with the floor at any point in time (Bruijn and van Dieen, 2018). In anteroposterior direction, the body CoM moves outside of the BoS during each of the single support phases of the gait cycle, which poses a challenge to stabilizing gait (Shimba, 1984, Winter, 1995).

The most extensively studied mechanism to stabilize gait is foot placement (Bauby and Kuo, 2000, Townsend, 1985, Wang and Srinivasan, 2014, Vlutters et al., 2016). Foot placement is considered the main mechanism for stabilizing gait in the AP movement direction (but also in mediolateral (ML) direction) (MacKinnon and Winter, 1993, Patla, 2003). A second mechanism is to apply active muscle moments around the ankle of the stance foot (‘ankle strategy’) (Horak and Nashner, 1986). Experimental data showed that humans adjust sagittal plane muscle moments around the ankle of the stance foot following AP mechanical perturbations of gait (Vlutters et al., 2016) (but also frontal plane muscle moments following ML perturbations (Hof and Duysens, 2018)). These ankle moments are reflected in a shift of the center of pressure of the ground reaction force (CoP). Whereas foot placement primarily moves the BoS to accommodate the state of the CoM, it also determines the location of the CoP. In both cases, CoP shifts cause acceleration of the CoM, allowing control of the CoM relative to the BoS.

An additional mechanism that can be used to stabilize gait, is the counter-rotation mechanism, i.e., changing the angular momentum of segments around the CoM to change the direction of the ground reaction force (Hof, 2007). In this mechanism, body segments are rotated with respect to the CoM (Otten, 1999). A forward acceleration of the trunk towards flexion for example results in a backward acceleration of the CoM and vice versa (Hof et al., 2007, Otten, 1999, Hof, 2007). This is often coined the hip strategy, but rotations of other body segments, for example arms or legs can be used in the same way. The appropriate regulation of whole-body angular momentum is essential for maintaining stable gait (Herr and Popovic, 2008). Whole-body angular momentum has been used to investigate how younger and older healthy adults stabilize the gait pattern over a range of walking tasks such as steady-state walking (Herr and Popovic, 2008), walking at different speeds (Bennett et al., 2010, Thielemans et al., 2014), walking at different step lengths (Thielemans et al., 2014), walking with an additional weight on the wrist or ankle (Thielemans et al., 2014), incline/decline walking (Silverman et al., 2012), stair ascent/descent (Silverman et al., 2014), and recovering from a trip (Pijnappels et al., 2005, Pijnappels et al., 2004, Potocanac et al., 2014). These studies have shown that the range of angular momentum during walking is kept low through the cancellation of angular momenta between body segments. However, the range of whole-body angular momentum has been found to increase when stable gait is compromised in the presence of perturbations (Martelli et al., 2013, Sheehan et al., 2015).

In healthy young adults, shifts of the CoP through modulation of foot placement and ankle moments (CoP-mechanism) appear to be predominantly used to stabilize gait, but the use of the counter-rotation mechanism likely increases with the difficulty of the task or the intensity of perturbations (Horak, 2006). Understanding the mechanisms used to stabilize gait may have implications for the design of therapeutic interventions that aim to decrease fall incidence. As stability in AP direction is challenged during walking due to movements of the body CoM outside of the BoS, the overall goal of this study was to determine the relative contribution of the CoP-mechanism and the counter-rotation mechanism to control the CoM in the AP direction during a normal step and the first recovery step after perturbation in healthy adults.

## Methods

### Subjects

Nineteen healthy volunteers (10 males 9 females, age 24.4±1.8 years old, weight 70.3 ±10.2 kg, height 1.75±0.08 m, means±s.d.) participated in this study. Exclusion criteria were surgery and current injury to lower extremities. The study was approved by the Institutional Review Board of the Faculty of Human Movement Sciences, VU University, Amsterdam, The Netherlands (ECB/2013-60). Before participating, every subject signed an informed consent form.

### Materials

The study took place in a Gait Real-time Analysis Interactive Lab (GRAIL; Motek, Amsterdam, the Netherlands). The GRAIL includes a split-belt treadmill with two integrated force plates (sample rate was 1000 samples per second), and a 3D motion capture system (Vicon Motion Systems, Oxford, United Kingdom) (sample rate was 100 samples per second). The two belts of the treadmill can be accelerated individually (e.g. to simulate slips). The Human Body Model (Motek, Amsterdam, The Netherlands) containing 55 anatomical markers was used to obtain full body kinematics (van den Bogert et al., 2013). Acceleration perturbations were timed by the software controlling the GRAIL (D-Flow; Motek, Amsterdam, the Netherlands). Even though the perturbations were not intended to make the subjects fall, a safety harness connected to the ceiling was worn.

### Research design

This study was part of a larger project focusing on validity measures for walking stability containing different perturbations types (accelerations, decelerations and medial and lateral sway) and 5 different perturbations magnitudes applied during the stance phase of walking (ECB/2013-60).

Before the study started, subjects were provided five minutes to get familiar with walking on the treadmill and with the lowest and highest perturbation magnitudes. In this study, subjects walked on the treadmill at fixed speed (1.2ms^−1^), while wearing comfortable flat-soled shoes. Unexpected perturbations consisting of belt accelerations were applied immediately after right heel-strike, which were determined in the D-flow software based on heel and sacrum markers (Zeni et al., 2008). Subjects were exposed to five different perturbation magnitudes, with varying speed differences relative to the fixed gait speed ranging from 0.1 and 0.5 m s^−1^ in steps of 0.1 m s^−1^ (P1-P5), designed to be finished within the stance phase of the right foot (Fig. 1).

**Fig. 1.**
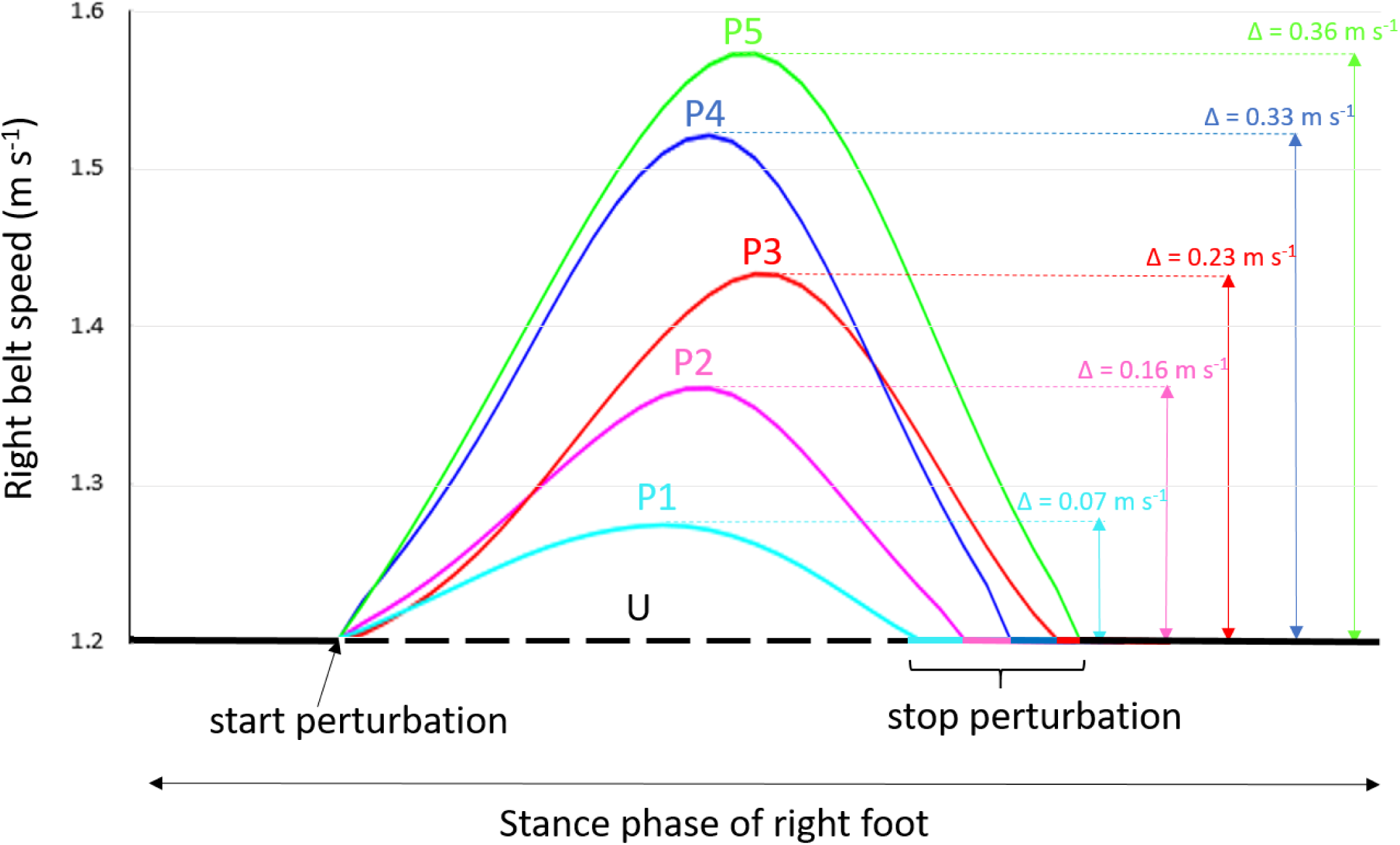
Illustration of the perturbations (split-belt accelerations). The arrows indicate the maximum right belt speed difference (Δ) between an unperturbed step (U) and the first recovery step after perturbation (P1-P5) during the stance phase of the right foot. The colors indicate the various perturbation magnitudes (P1-P5).

Subjects performed 5 trials, each of which contained 15 perturbations of which magnitude and number of steps between subsequent perturbations were randomly varied. This resulted in 15 repetitions at each perturbation magnitude. Time interval between subsequent perturbations was varied between 10 and 15 strides.

### Data analysis

Kinematic and force plate data were low-pass filtered at 6Hz using a bi-directional fourth order Butterworth filter. Kinematic data were analyzed using a 16-segment kinematic model. For each segment, mass, CoM, origin, and inertia tensor were calculated as described previously (Zatsiorski, 1998, Kingma et al., 1996, Faber et al., 2011). Full body CoM was calculated by combining the CoM of all segments. Heel-strikes and toe-offs were determined based on CoP data (Roerdink et al., 2008).

The contributions of the CoP-mechanism and counter-rotation mechanism to the CoM acceleration, as described by Hof (Hof, 2007) were calculated for the AP direction using Eqn 1:

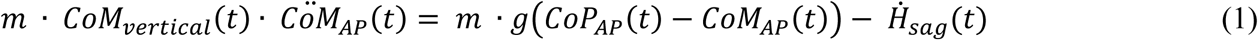

in which *m* is body mass, *CoM*_*vertical*_ and *CoM*_*AP*_ are the vertical and AP position of the CoM, *CÖM*_*AP*_ is the double derivative of *CoM*_*AP*_, *t* is time, *g* is the gravitational acceleration, *CoP*_*AP*_ is the AP position of the CoP, and 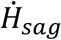 is the change in total body angular momentum in the sagittal plane.

Here, the first part of the right-hand term can be seen as the AP CoM acceleration due to displacement of the CoP (i.e. the contribution of ankle moments and foot placement to CoM acceleration), and 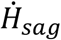, the second part, is proportional to the AP CoM acceleration induced by angular momentum change (i.e. the contribution of the counter-rotation mechanism to the CoM acceleration).

The contribution of the CoP and the counter-rotation mechanism to CoM acceleration were calculated during an unperturbed step and during the first recovery step after each perturbation (P1-P5). Analysis started at left heel-strike, i.e., after the belt reached constant speed again, to avoid that horizontal forces associated with belt acceleration would affect the mechanical analysis. For description of the results, the (left) step was divided into double support phase 1 (from left heel-strike until right toe-off), left single leg stance phase 1 (from right toe-off until mid-stance), left single leg stance phase 2 (from mid-stance until right heel-strike), and double support phase 2 (from right heel-strike until left toe-off) (Fig. 2).

**Fig. 2.**
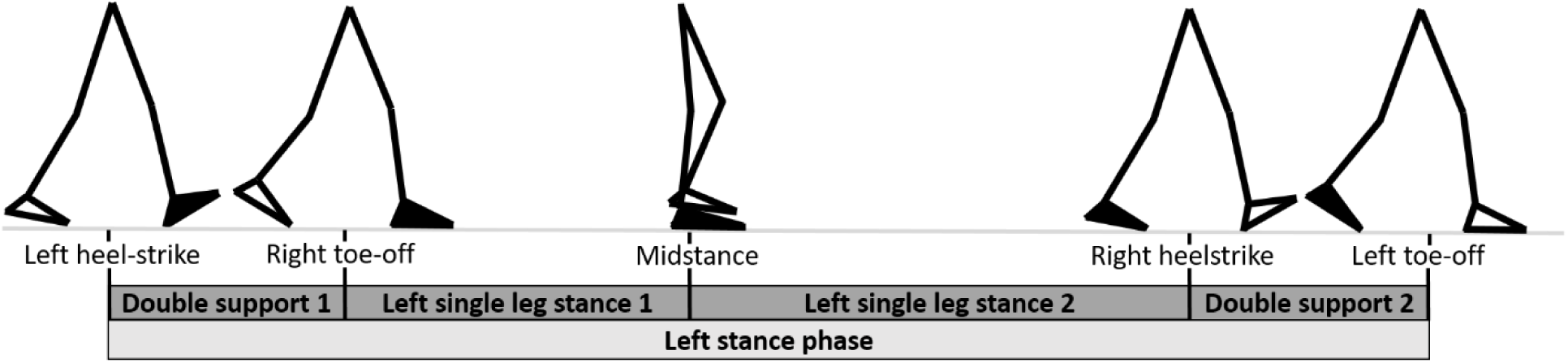
Gait events during the left stance phase with the left leg dominant (black foot).

### Statistics

Non-parametric statistical tests were used, as the D’Agostino-Pearson K2 test revealed that the values were not normally distributed.

SPM non-parametric repeated measures ANOVA was performed using open-source SPM1d code (vM.0.4.5, www.spm1D.org), comparing the CoM acceleration and the contributions of the two mechanisms (CoP and counter-rotation mechanism) to the CoM acceleration in the AP direction between an unperturbed step and during the first recovery step after the perturbation (α = 0.05).

First, we compared the time normalized AP CoM accelerations between the conditions. Next, we determined which mechanisms were used to control the CoM, by assessing differences in time normalized curves of the CoP and counter-rotation mechanism between the conditions.

For each SPM repeated measures ANOVA, a statistical parametric map (SPM(F) or SPM(t) respectively) was created by calculating the conventional univariate t- or F-statistic at each point of the gait cycle (Pataky, 2010). Afterwards, Random Field Theory allowed an estimation of the critical threshold that only 5% (α = 0.05) of equally smooth random data are expected to exceed (Adler and Taylor, 2007). If the SPM(F) crossed the critical threshold, indicating a significant main effect, post-hoc SPM(t) maps were calculated for within-group comparisons and a supra-threshold cluster was created, indicating a significant difference between the two conditions in a specific phase of the gait cycle.

A Bonferroni correction was applied for each comparison of the AP CoM acceleration, the contribution of the CoP-mechanism and the contribution of the counter-rotation mechanism between six conditions (unperturbed step and the first recovery step after P1-P5), to adjust α for multiple post-hoc comparisons (α = 0.05/15 = 0.003). P-values < 0.003 were considered statistically significant for the multiple post-hoc comparisons (Altman, 1991).

## Results

Fig. 3 shows the CoM acceleration and the contribution of the CoP-mechanism and the counter-rotation mechanism to the CoM acceleration during a normal unperturbed step and during the first recovery step after perturbations with different magnitudes.

**Fig. 3.**
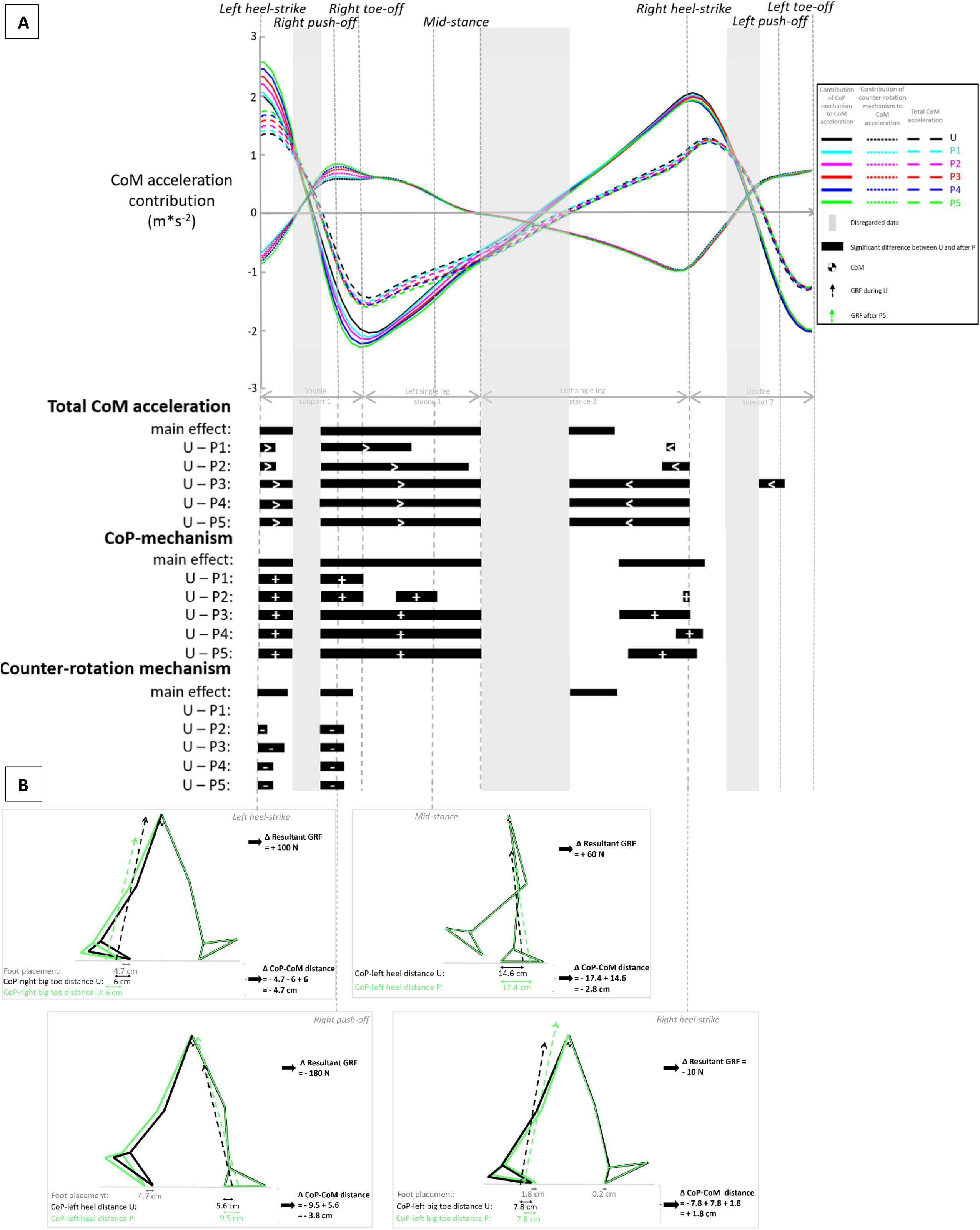
***3A;*** Averaged time series (N=19) of the center of mass (CoM) acceleration (dashed lines) and the contribution of the center of pressure mechanism (CoP-mechanism) (solid lines) and the counter-rotation mechanism (dotted lines) to the CoM acceleration during a unperturbed step (U) and during the first recovery step after perturbations with different magnitudes (P1-P5). Colors indicate the perturbation magnitudes. Grey bars indicate disregarded data where the CoM acceleration approaches zero. Black bars indicate gait phases with significant differences between U and the first recovery step after P (main effect; p<0.05, post-hoc comparisons; p<0.003). The greater then sign (>) indicates that the total CoM acceleration is greater during the first recovery step after P compared to U. The less then sign (<) indicates that the total CoM acceleration is smaller during the first recovery step after P compared to U. The plus sign (+) indicates that the mechanism contributes to the total CoM acceleration. The minus sign (-) indicates that the mechanism counteracts the total CoM acceleration. ***3B;*** Stick Figures of the lower body of a typical subject at left heel-strike, right push-off, mid-stance and right heel-strike including the Ground Reaction Force (GRF) during U (black) and after P5 (green).

During the first half of double support phase 1, the CoM accelerated forward and more so in perturbed than unperturbed gait, reflecting the tendency to fall forward due to the belt acceleration. The backward shift of the CoP, induced by the belt acceleration, contributed to this increased CoM acceleration, while a negative contribution of the counter-rotation counteracted this increased CoM acceleration (Fig. 3A). Late in this phase, from right push-off until mid-stance, the CoM decelerated both in unperturbed and perturbed walking, but this deceleration was larger in perturbed walking indicative of balance recovery. The CoP mechanism contributed to the deceleration as indicated by the larger negative contribution, while the counter-rotation mechanism contributed a larger positive acceleration and thus amplified the effect of the perturbation on the CoM. The effects of the perturbations on total CoM acceleration and on the contribution of the CoP mechanism remained visible throughout left single stance 1, while the counter-rotation mechanism was no longer different from normal walking from single stance onwards. During left single leg stance phase 2 and double support phase 2, CoM acceleration was less positive in perturbed than unperturbed walking, indicating continued correction of the effect of the perturbation. The CoP-mechanism contributed to reducing the CoM acceleration as indicated by the less positive values in perturbed compared to unperturbed steps (Fig. 3A). These differences between the first recovery step after a perturbation and an unperturbed step were present even after the perturbations with the lowest magnitudes. However, effects were not observed for the same time intervals for all perturbation magnitudes (Fig. 3A).

### CoP shifts determining the contribution of the CoP-mechanism to the CoM acceleration

Fig. 3B shows the body configuration of a typical subject and the ground reaction force vector during an unperturbed step and during a step after perturbation P5 to illustrate how the CoP-mechanism contributed to recovery.

#### Left heel-strike

At left heel-strike, as a result of the belt acceleration during right stance, the distance between the right foot and the CoM was 4.7 cm greater after perturbation P5 compared to an unperturbed step (Fig. 3B). The CoP location within the right foot was comparable between a perturbed and unperturbed step. This resulted in a contribution of the CoP-mechanism to the larger positive CoM acceleration after perturbation (Fig. 3A).

#### Right push-off

The CoP location relative to the left heel during right push-off was 3.8 cm greater after perturbation P5 compared to an unperturbed step, which increased the distance between the CoP and the CoM by 3.8 cm after P5 compared to an unperturbed step (Fig. 3B). This resulted in a more negative CoM acceleration after the perturbation (Fig. 3A).

#### Mid-stance

The CoP location relative to the left heel was 2.8 cm greater after perturbation P5 compared to an unperturbed step, which increased the distance between the CoP and the CoM by 2.8 cm after P5 compared to an unperturbed step (Fig. 3B). This resulted in a more negative CoM acceleration after the perturbation (Fig. 3A).

#### Right heel-strike

At right heel-strike, the distance between the left foot and the CoM was 1.8 cm smaller after perturbation P5 than in an unperturbed step (Fig. 3B). The CoP location within the left foot was comparable between a perturbed and unperturbed step. (Fig. 3B). Consequently, the distance between the CoP and the CoM was 1.8 cm smaller after perturbation P5 compared to an unperturbed step. This resulted in a lower positive contribution of the CoP-mechanism, contributing to the decrease of the positive CoM acceleration after the perturbation (Fig. 3A).

## Discussion

The goal of this study was to determine the relative contribution of the CoP-mechanism and the counter-rotation mechanism to the control of the CoM in the AP direction in recovering from ‘trip-like’ perturbations during walking in healthy adults. We found that the CoP-mechanism contributed to corrections of the CoM acceleration after perturbations in the AP direction, while the counter-rotation mechanism actually contributed to CoM acceleration in the direction of the perturbation, but only in the initial phases of the first step after the perturbation. Interestingly, the CoP and counter-rotation mechanisms counteracted each other consistently in both unperturbed and perturbed gait.

### Comparison unperturbed and perturbed step

The CoM acceleration was significantly more positive during the first part of the double support phase 1 after the perturbation. This is probably a direct effect of the belt acceleration which shifts the right foot backward relative to the body causing a forward directed moment of the ground reaction force. The decreased ground reaction force after the perturbation at this point in time may suggest an initial inhibitory response to the perturbation. The CoM acceleration was more negative after perturbations from right push-off onwards indicating a corrective response. This corrective response was driven by the CoP-mechanism, while the counter-rotation mechanism actually worked in opposite direction, enhancing the effect of the perturbation.

Previous work showed that in the AP direction, ankle moments are key in adjusting the CoP location, and therefore in regulating the body’s accelerations (Vlutters et al., 2016, Gruben and Boehm, 2014). This is in line with the results of the current study. Vlutters et al., (2016) found that healthy subjects did not significantly adjust their foot placement relative to the CoM in the first step following a pelvis perturbation in AP direction applied at toe-off (Vlutters et al., 2016). However, differentiating between the ankle moments and foot placement to induce CoM accelerations was difficult in this study, because the perturbation (a belt acceleration) pulled the right foot backwards, most likely also affecting left foot placement. This is why effects of foot placement and ankle moments were merged into the CoP-mechanism in this study.

The increased contribution of the counter-rotation mechanism to CoM acceleration around left heel-strike and right push-off after perturbations P2-P5 is in line with findings of Sheenan et al., (2015), who found that the range of whole-body angular momentum increases when perturbations are present (Sheehan et al., 2015). However, the counter-rotation mechanism counteracted the desired CoM acceleration. Vlutters et al., (2018) reported that, while experiencing AP perturbations during walking with ineffective ankles by using pin-shoes, subjects did not use a hip strategy, but relied on foot placement adjustments instead (Vlutters et al., 2018). This suggests a low priority for the hip strategy, which could be because the hip strategy would interfere with the gait pattern. Altering the angular momentum in the sagittal plane will strongly affect the leg swing and modifying this will obviously induce inappropriate foot placement (Vlutters et al., 2018, Oddsson et al., 2004). Our results confirm this, as the contribution of the counter-rotation mechanism to the CoM acceleration counteracted the desired CoM acceleration in the initial phases of the first step after the perturbation and did not differ between an unperturbed step and a perturbed step during later phases of the first step after perturbation

### Limitations of the current study

Perturbation magnitudes were lower than intended. At the highest magnitude a belt speed difference of 0.36 m s^−1^ was reached, instead of the intended 0.5 m s^−1^. However, the perturbation magnitudes in this study were high enough to consistently elicit significant differences in the contribution of the mechanisms to the CoM acceleration between perturbed and unperturbed walking. Nevertheless, differentiating between the ankle moments and foot placement to induce these CoM accelerations was difficult in this study, because the perturbation (a belt acceleration) pulled the right foot backwards, most likely also affecting left foot placement.

Treadmill walking aims to simulate overground walking, but treadmill walking imposes various constraints on the subject that are not present during overground walking. The treadmill width is limited (~1 m), but most importantly the treadmill requires the subject to continue walking. The latter may constrain recovery responses as responses leading to a complete stop would be undesirable. However, a comparison of joint kinematics and ground reaction forces between treadmill and overground walking conditions suggests that differences between the two conditions are within the normal variability of gait at a given speed (Riley et al., 2007). Furthermore, in a study by Zadravec et al., (2017) two similar perturbation devices were used to compare human stepping in response to pelvis perturbations during both treadmill and overground walking conditions (Zadravec et al., 2017). They concluded that the responses in both conditions were similar.

## Conclusions

We found that the CoP-mechanism contributed to corrections of the CoM acceleration after perturbations in the AP direction, while the counter-rotation mechanism actually counteracted the desired CoM acceleration, but only in the initial phases of the first step after the perturbation. Interestingly, the CoP and counter-rotation mechanisms counteracted each other consistently in both unperturbed and perturbed gait. The CoP-mechanism regulated the CoM acceleration after perturbation in the AP direction. Whereas the counter-rotation mechanism appeared to prevent interference with the gait pattern, rather than using it to influence the CoM acceleration after the perturbation in AP direction. This is the case, because the angular moment in the sagittal plane is strongly affected by leg swing and modifying this obviously has consequence for appropriate foot placement.

## List of symbols and abbreviations

CoM: center of mass
BoS: base of support
AP: anteroposterior
ML: mediolateral
CoP: center of pressure
GRAIL: Gait Real-time Analysis Interactive Lab
P1-P5: perturbation magnitudes (P1; lowest magnitude, P5: highest magnitude)
*m*: body mass
CoM_vertical_: vertical position of the CoM
CoM_AP_: AP position of the CoM
*t*: time
CöM_AP_: double derivative of CoM_AP_
*g*: gravitational acceleration
CoP_AP_: AP position of the CoP
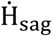: change in total body angular momentum in the sagittal plane

## Acknowledgments

The authors gratefully acknowledge Lars Aarts and Josien Rozendaal for their help during the experiments and Sanne Roeles from Motekforce Link for technical support.

## Data availability

The data that support the findings of this study are available on request from the corresponding author, [MvdB].

## References

Adler, R. J. & Taylor, J. E. 2007. Random Fields and Geometry. New York: Springer-Verlag 1st edition.

Altman, D. G. 1991. Practical statistics for medical research. London: Chapman and Hall.

Bauby, C. E. & Kuo, A. D. 2000. Active control of lateral balance in human walking. J Biomech, 33, 1433–40.

Bennett, B. C., Russell, S. D., Sheth, P. & Abel, M. F. 2010. Angular momentum of walking at different speeds. Hum Mov Sci, 29, 114–24.

Bruijn, S. M., Meijer, O. G., Beek, P. J. & Van Dieen, J. H. 2013. Assessing the stability of human locomotion: a review of current measures. J R Soc Interface, 10, 20120999.

Bruijn, S. M. & Van Dieen, J. H. 2018. Control of human gait stability through foot placement. J R Soc Interface, 15.

Faber, G. S., Kingma, I. & Van Dieen, J. H. 2011. Effect of initial horizontal object position on peak L5/S1 moments in manual lifting is dependent on task type and familiarity with alternative lifting strategies. Ergonomics, 54, 72–81.

Gruben, K. G. & Boehm, W. L. 2014. Ankle torque control that shifts the center of pressure from heel to toe contributes non-zero sagittal plane angular momentum during human walking. J Biomech, 47, 1389–94.

Herr, H. & Popovic, M. 2008. Angular momentum in human walking. J Exp Biol, 211, 467–81.

Hof, A. L. 2007. The equations of motion for a standing human reveal three mechanisms for balance. J Biomech, 40, 451–7.

Hof, A. L. & Duysens, J. 2018. Responses of human ankle muscles to mediolateral balance perturbations during walking. Hum Mov Sci, 57, 69–82.

Hof, A. L., Van Bockel, R. M., Schoppen, T. & Postema, K. 2007. Control of lateral balance in walking. Experimental findings in normal subjects and above-knee amputees. Gait Posture, 25, 250–8.

Horak, F. B. 2006. Postural orientation and equilibrium: what do we need to know about neural control of balance to prevent falls? Age Ageing, 35 Suppl 2, ii7–ii11.

Horak, F. B. & Nashner, L. M. 1986. Central programming of postural movements: adaptation to altered support-surface configurations. J Neurophysiol, 55, 1369–81.

Kingma, I., Delooze, M. P., Toussaint, H. M., Klijnsma, H. G. & Bruijnen, T. B. M. 1996. Validation of a full body 3-D dynamic linked segment model. Human Movement Science, 15, 833–860.

Mackinnon, C. D. & Winter, D. A. 1993. Control of whole body balance in the frontal plane during human walking. J Biomech, 26, 633–44.

Martelli, D., Monaco, V., Bassi Luciani, L. & Micera, S. 2013. Angular momentum during unexpected multidirectional perturbations delivered while walking. IEEE Trans Biomed Eng, 60, 1785–95.

Oddsson, L. I., Wall, C., Mcpartland, M. D., Krebs, D. E. & Tucker, C. A. 2004. Recovery from perturbations during paced walking. Gait Posture, 19, 24–34.

Otten, E. 1999. Balancing on a narrow ridge: biomechanics and control. Philos Trans R Soc Lond B Biol Sci, 354, 869–75.

Pataky, T. C. 2010. Generalized n-dimensional biomechanical field analysis using statistical parametric mapping. J Biomech, 43, 1976–82.

Patla, A. E. 2003. Strategies for dynamic stability during adaptive human locomotion. IEEE Eng Med Biol Mag, 22, 48–52.

Pijnappels, M., Bobbert, M. F. & Van Dieen, J. H. 2004. Contribution of the support limb in control of angular momentum after tripping. J Biomech, 37, 1811–8.

Pijnappels, M., Bobbert, M. F. & Van Dieen, J. H. 2005. Push-off reactions in recovery after tripping discriminate young subjects, older non-fallers and older fallers. Gait Posture, 21, 388–94.

Potocanac, Z., De Bruin, J., Van Der Veen, S., Verschueren, S., Van Dieen, J., Duysens, J. & Pijnappels, M. 2014. Fast online corrections of tripping responses. Exp Brain Res, 232, 3579–90.

Riley, P. O., Paolini, G., Della Croce, U., Paylo, K. W. & Kerrigan, D. C. 2007. A kinematic and kinetic comparison of overground and treadmill walking in healthy subjects. Gait Posture, 26, 17–24.

Roerdink, M., Coolen, B. H., Clairbois, B. H., Lamoth, C. J. & Beek, P. J. 2008. Online gait event detection using a large force platform embedded in a treadmill. J Biomech, 41, 2628–32.

Sheehan, R. C., Beltran, E. J., Dingwell, J. B. & Wilken, J. M. 2015. Mediolateral angular momentum changes in persons with amputation during perturbed walking. Gait Posture, 41, 795–800.

Shimba, T. 1984. An estimation of center of gravity from force platform data. J Biomech, 17, 53–60.

Silverman, A. K., Neptune, R. R., Sinitski, E. H. & Wilken, J. M. 2014. Whole-body angular momentum during stair ascent and descent. Gait Posture, 39, 1109–14.

Silverman, A. K., Wilken, J. M., Sinitski, E. H. & Neptune, R. R. 2012. Whole-body angular momentum in incline and decline walking. J Biomech, 45, 965–71.

Thielemans, V., Meyns, P. & Bruijn, S. M. 2014. Is angular momentum in the horizontal plane during gait a controlled variable? Hum Mov Sci, 34, 205–16.

Townsend, M. A. 1985. Biped gait stabilization via foot placement. J Biomech, 18, 21–38.

Van Den Bogert, A. J., Geijtenbeek, T., Even-Zohar, O., Steenbrink, F. & Hardin, E. C. 2013. A real-time system for biomechanical analysis of human movement and muscle function. Med Biol Eng Comput, 51, 1069–77.

Vlutters, M., Van Asseldonk, E. H. & Van Der Kooij, H. 2016. Center of mass velocity-based predictions in balance recovery following pelvis perturbations during human walking. J Exp Biol, 219, 1514–23.

Vlutters, M., Van Asseldonk, E. H. F. & Van Der Kooij, H. 2018. Reduced center of pressure modulation elicits foot placement adjustments, but no additional trunk motion during anteroposterior-perturbed walking. J Biomech, 68, 93–98.

Wang, Y. & Srinivasan, M. 2014. Stepping in the direction of the fall: the next foot placement can be predicted from current upper body state in steady-state walking. Biol Lett, 10.

Winter, D. A. 1995. Human balance and posture control during standing and walking. Gait Posture 3, 193–214.

Zadravec, M., Olensek, A. & Matjacic, Z. 2017. The comparison of stepping responses following perturbations applied to pelvis during overground and treadmill walking. Technol Health Care, 25, 781–790.

Zatsiorski, V. M. 1998. Kinematics of human motion, Champaign IL, Human Kinetics.

Zeni, J. A., Jr., Richards, J. G. & Higginson, J. S. 2008. Two simple methods for determining gait events during treadmill and overground walking using kinematic data. Gait Posture, 27, 710–4.

